# Statistical Coupling Analysis Predicts Correlated Motions in Dihydrofolate Reductase

**DOI:** 10.1101/2024.06.18.599103

**Authors:** Thomas L. Kalmer, Christine Mae F. Ancajas, Cameron I. Cohen, Jade M. McDaniel, Abiodun S. Oyedele, Hannah L. Thirman, Allison S. Walker

## Abstract

The role of dynamics in enzymatic function is a highly debated topic. Dihydrofolate reductase (DHFR), due to its universality and the depth with which it has been studied, is a model system in this debate. Myriad previous works have identified networks of residues in positions near to and remote from the active site that are involved in dynamics and others that are important for catalysis. For example, specific mutations on the Met20 loop in *E. coli* DHFR (N23PP/S148A) are known to disrupt millisecond-timescale motions and reduce catalytic activity. However, how and if networks of dynamically coupled residues influence the evolution of DHFR is still an unanswered question. In this study, we first identify, by statistical coupling analysis and molecular dynamic simulations, a network of coevolving residues, which possess increased correlated motions. We then go on to show that allosteric communication in this network is selectively knocked down in N23PP/S148A mutant *E. coli* DHFR. Finally, we identify two sites in the human DHFR sector which may accommodate the Met20 loop double proline mutation while preserving dynamics. These findings strongly implicate protein dynamics as a driving force for evolution.

## Introduction

Enzymes play a crucial role in nearly all biological processes, catalyzing reactions that would otherwise be inaccessible in nature. The mechanism of these enzymes can vary greatly, but they all function to accelerate chemical reactions by lowering activation energy. In this pursuit, enzymes move in a variety of ways to form the interactions necessary for catalysis. However, the field has yet to come to a consensus about just how much these movements, both large and small, contribute to the catalytic power of enzymes.^1–4^ Many publications have covered this debate, with several suggesting that much of the controversy is caused by the lack of clarity in important definitions, such as what timescale motions should occur in, how to define contribution to enzymatic power and what experiments should be performed in order to prove significance.^1,2,4^

With the introduction and expansion of molecular dynamics (MD) simulations, along with other movement-encompassing techniques, an increasing number of results from biochemical literature suggest that dynamical motions, fundamental for catalytic power or not, are involved in a wide variety of enzymatic functions.^4–6^ These studies provide crucial insights for their respective systems. Moreover, since reaction rate differences of just one order of magnitude can easily impact survival, very small differences in enzymatic function as a consequence of hindered dynamics can impose considerable evolutionary pressure on biological systems.^2^ For this reason, it is logical to think that evolution may act to conserve specific amino acids or networks of amino acids as a mechanism for preserving important motions.

A model system to study the role of protein dynamics is dihydrofolate reductase (DHFR). DHFR is an enzyme responsible for the reduction of dihydrofolate to tetrahydrofolate in which the cofactor nicotinamide adenine dinucleotide phosphate (NADPH) acts as a hydride donor.^7–10^ Tetrahydrofolate and derivatives are essential for thymidylate and purine synthesis and maintenance of reduced folate pools.^11,12^ Subsequently, inhibition of DHFR results in a disruption of DNA replication and eventual cell death.^11,13^ Due to its critical role in cell health, DHFR has become an attractive drug target for multiple diseases.^14,15^ Many DHFR inhibitors are already available to treat a wide range of diseases including fungal infections, parasites, cancer, arthritis, Crohn’s disease, and many other inflammatory conditions.^12,16,17^ DHFR is also highly structurally conserved across the tree of life, although sequence homology is weak.^10,18–20^ Multiple studies have been conducted on the kinetics and conformational changes of DHFR, especially in *Escherichia coli* (*ec*DHFR).^7,8,21^ DHFR relies on a series of ligand-induced conformational changes to facilitate catalysis which in humans (*h*DHFR) occurs through a hinge opening motion of the enzyme.^5,19,22–24^ The enzyme exists in the hinge-open conformation when empty and then changes to the hinge-closed conformation upon ligand binding. The latter conformation tightly packs the active site and favors hydride transfer from NADPH.

Meanwhile, the Met20 loop in *ec*DHFR is more flexible and adapts distinct occluded and closed conformations throughout the enzymatic cycle.^19^ Notably, though the human and *ec*DHFR undergo the same catalytic reaction, they differ in their catalytic efficiency and dynamic behavior.^18^ Despite extensive studies on DHFR, further studies are required to assess the contribution of dynamics to catalysis and the role of evolution.^4–6^

Dynamic coupling in proteins is a mechanism in which two sites on a protein molecule are dynamically linked, enabling long-range communication and cooperative interactions, despite not necessarily being in direct physical contact.^25^ This mechanism is a crucial aspect of allosteric regulation, whereby ligand binding or mutation-induced conformational changes trigger responses in remote regions of the protein.^26^ The study of protein dynamics provides an understanding of biological processes at a molecular level, including signal transmission, protein interactions, disordered protein behavior, and nucleic acid movements.^6,27^ Statistical Coupling Analysis (SCA), on the other hand, is a computational method that has been developed to analyze multiple sequence alignments (MSAs) of protein families in order to identify groups of coevolving residues that are referred to as “sectors”.^28,29^ These sectors represent spatially organized networks within protein structures that often connect positions in the active site to surface sites distributed throughout the protein. ^25^ The application of SCA serves as a valuable tool for researchers to identify functionally critical sectors within proteins and shed light on the propagation and dissipation of perturbations within protein structures^.30^ SCA has been instrumental in identifying networks of coevolved amino acids in proteins, such as in the MutS DNA mismatch repair protein, explaining the allosteric regulation and protein dynamics.^30–32^ This method allows for the quantitative examination of the long-term correlated evolution of amino acids within protein families, highlighting the statistical signature of functional constraints arising from conserved communication between positions.^33^

While SCA has previously been used to successfully identify allosteric networks within a variety of proteins,^34–39^ the physicochemical interactions that enable communication through these allosteric networks and how evolutionary constraints are defined by these interactions are still poorly understood.^40–42^ The few studies that have examined how SCA sectors relate to dynamics have mostly focused on using SCA to predict which mutations will impact dynamics^31^ or combining SCA and molecular dynamics simulations to identify coupled positions^32^ or allosteric pockets for drug design.^43^ There have been relatively few studies that investigated how coevolving residues identified by SCA relate to dynamic networks within proteins, although the theoretical underpinnings of SCA and molecular evolution suggest that dynamic networks that are important for protein function should be represented as sectors. One study did find a strong overlap between sectors identified by a decomposition of a covariance matrix of structural dynamics and the sectors identified by SCA.^32^ However, this study was limited to a PDZ domain, which primarily serve as anchoring domains and do not have enzymatic activity. Therefore, it is unclear if the same relationship between SCA sectors and dynamic networks would be present in enzymes such as DHFR. Further investigation is required to understand the relationship between evolutionary and dynamic networks.

Here, we investigate how correlated motions in DHFR relate to coevolution of residues. To assess this, we utilized SCA and ran MD simulations on wild-type *ec*DHFR. SCA identified sectors and independent components containing coevolving residues throughout the enzyme, and pairwise dynamic cross correlation was used to detect dynamic motions. We further analyze how perturbations of residues interacting in these SCA-identified networks affect dynamics by performing simulations on a previously designed N23PP/S148A mutant *ec*DHFR. Our findings illustrated decreased dynamics in the mutant *ec*DHFR. Furthermore, we discussed how evolutionary constraints may relate to protein dynamics and identified specific changes in the human DHFR sector relative to *E. coli*.

## Methods

### Statistical Coupling Analysis (SCA)

A representative protein alignment of 4422 sequences with 802 positions of DHFR was obtained from PFAM (PF00186) using the Stockholm 1.0 format. Statistical coupling analysis was performed on the sequence alignment using the python package pySCA6.0.^37^ After processing, the final alignment size was 3664 sequences with 146 positions, and following calculation of sequence weights there were 3216 effective sequences. The scaSectorID script identified four groups of coevolving residues, or independent components (IC), in DHFR. After fitting to an empirical statistical distribution, a cutoff of p=0.95 was used to determine positions which significantly contributed to each IC. The full pySCA package is available on the Ranganathan lab github (https://github.com/ranganathanlab). Additionally, residues in each IC were ordered numerically and an SCA by IC matrix was constructed. Code available on github. (https://github.com/Kalmertl/Statistical-Coupling-Analysis-Predicts-Correlated-Motions-in-Dihydrofolate-Reductase)

### Molecular Dynamics (MD) Simulation

The MD simulations were conducted using AmberTools22 and Amber22 suite.^44,45^ Protein Data Bank^46^ entries 3QL3 and 3QL0 were used as the initial structures for the wild-type and mutant DHFRs, respectively.^47^ Nicotinamide adenine dinucleotide phosphate (NAP) cofactor and folate (FOL) ligand geometry were extracted from the appropriate PDB entry. FOL was protonated in GaussView and parameterized with Antechamber with the following options “-c bcc -nc -2 -at gaff2”, and the NAP was protonated using reduce in LEaP, geometry optimized and energy minimized with Gaussian version 16 using the UFF molecular mechanics force field, and parameterized with Antechamber with the following options “-c bcc -nc -3 -at gaff2 -ek “scfconv=1.d-8 ndiis_attempts=1000 grms_tol=0.002”. H++^48^ was used to obtain the correct protonation states of the amino acids throughout the structures at a pH of 6.5. The program *xleap*^*44*^ was used to apply ff19SB^49^ and GAFF2^50^ force fields to the proteins and ligands. The models were neutralized with Na^+^ ions and solvated using the SPC/E water model in a truncated octahedral box with a buffer of 14 Å. Before the MD simulation, a process of minimization, heating, and equilibration was performed. Steepest descent minimization of the solvated system with a restraining force of 500.0 kcal/mol Å^-2^ on the protein was initiated with 500 steps followed by 500 steps of conjugate gradient minimization. This process was iterated for the entire system at 0.0 kcal/mol Å^-2^ with 1000 steps of steepest descent minimization followed by another 1500 steps of conjugate gradient minimization. Subsequent equilibration and heating of the system from 0 K to 300 K using the Langevin temperature scheme and a 10.0 kcal/mol Å^-2^ constraint on the protein and ligands for over 20 ps simulation were performed. Final equilibration at constant pressure of 1 atm and constant temperature at 300 K for 100 ps was conducted to relax the system to an equilibrium density. Explicit solvent MD simulation was continued at constant pressure for each DHFR protein for 13 ns, and the time step was set to 2 fs with the trajectory snapshots saved at every 5 ps. The *cpptraj* package^51^ included in AmberTools22 was used to compute the dynamic cross correlation (DCC) matrix root-mean-squared deviations (RMSD). Molecular visualization was carried out using PyMOL.^52^

### DCC Comparison with SCA

Pairwise RMSD values from the DCC matrix were mapped according to their IC assignment. Any amino acid pairs within two positions of each other were excluded. Pairwise RMSD values where both residues fell within a given IC were assigned to that IC for analysis. Further, pairwise values were assigned to: “no IC” if neither amino acid in the pair belonged to an IC, “any IC” if both amino acids belonged to an IC but not necessarily the same IC, and “not within same IC” for all possible pairs except those within the same IC. Using the absolute values for pairwise correlations, significant differences between distributions were determined via a two-sided Mann-Whitney U test. DCC matrix residue numbering was shifted up by one after position 23 in the 3QL0 mutant to account for the insertion (position 23 became residue 24 etc.). RMSD value extraction, IC assignment, and statistical analysis were performed using a custom python script. GitHub available. (https://github.com/Kalmertl/Statistical-Coupling-Analysis-Predicts-Correlated-Motions-in-Dihydrofolate-Reductase)

## Results and Discussion

### SCA Identifies Four Independent Components in DHFR

To analyze the co-evolutionarily conserved residues within DHFR, we applied statistical coupling analysis (SCA).^28^ By examining the SCA matrix (**Figure 1A**), we identified coevolving residues, called sectors, encompassing the active site, cofactor- and substrate-binding sites, and other distant positions which are consistent with previous studies.^25,33,53^ As noted by Reynolds et al., several residues within these sectors coincide with millisecond fluctuations involved in important dynamic motions underlying catalysis.^25,47,54,55^ Additionally, other studies have underscored the importance of coevolving residues captured by SCA, linking them to essential protein functions.

**Figure 1.**
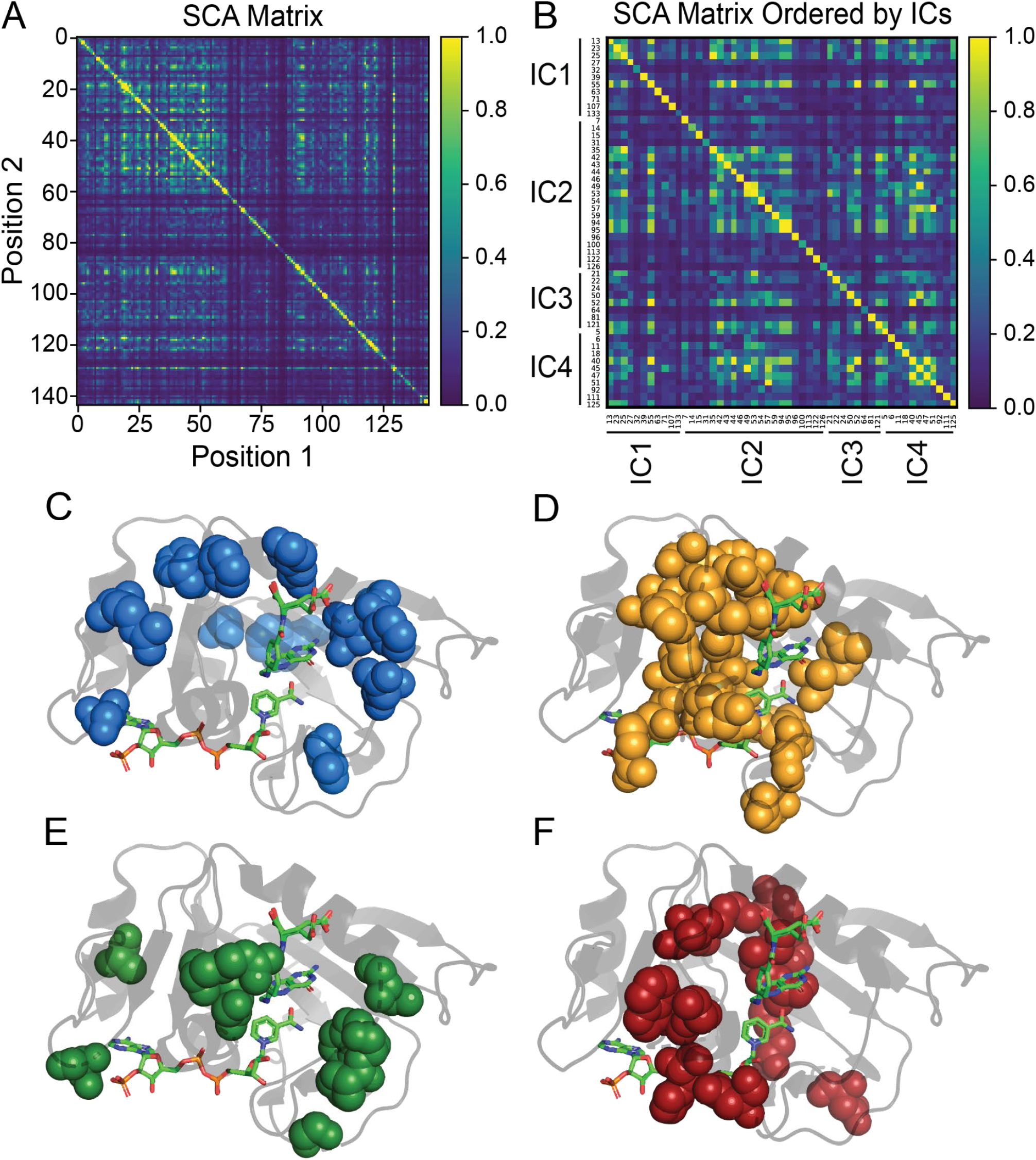
SCA matrix **(A)** constructed using the pySCA package as well as a custom matrix with IC contributors ordered numerically **(B)**. Additionally, the respective mapping of each independent component (IC1 - IC4) in DHFR as determined by statistical coupling analysis is shown **(C-F)**. The specific positions contributing to each IC can be found in panel B.

Through decomposition of the SCA matrix, we identified four coevolving independent components (ICs) within DHFR **(Figure 1B)**. Independent component analysis (ICA) is designed to maximize the statistical independence of ICs. In many cases ICs still show some dependence on each other, and these ICs can be combined into larger sectors.^37^ We observed dependence between ICs in DHFR (Figure 1B) but chose to analyze each IC separately to provide better insight into how ICs relate to protein dynamics. Mapping each of the ICs onto DHFR revealed ICs clustered around the active site as well as distantly positioned surface sites. IC1 and IC3 **(Figure 1C and 1E)** are spatially non-contiguous, while IC2 and IC4 **(Figure 1D and 1F)** demonstrated a high degree of physical connectivity, localizing mainly to the cofactor and substrate binding sites of DHFR. Previous studies have underscored the significant role many of the residues within these ICs play with regard to protein function and dynamics. Specifically, Asp27, found in the active site and captured by IC1, is known to coordinate with the pterin ring of folate.^56^ One functionally significant region within DHFR is the Met20 loop (residues 9-24) which changes conformation throughout the catalytic cycle of DHFR from the closed (E:NADPH and E:NADPH:DHF) to the occluded (E:NADP+:THF, E:THF, and E:NADPH:THF) states.^55^ Two other important loops are F-G (residues 116-132) and G-H (residues 142-150), providing stability with the Met20 loop.^10^ Several of these highlighted residues are captured within our four ICs. For example, Pro21, Trp22, and N23 which are involved in Met20 hinge motion are within IC-3. IC-4 also contains Tyr100 and Phe125 which were previously identified to play a key electrostatic role during hydride transfer while a theoretical mutagenesis study has also implicated Asp122 within IC2 to affect coupled functional motions.^56–60^ By further analyzing the SCA matrix, we begin to unravel the interactions within these networks of residues in the ICs. Whether these residues are near or distant from important sites in DHFR, their interactions suggest further implications in allosteric regulation and other functional dynamics.

### Molecular Dynamics Simulations of DHFR Reveal Correlated Motions Within ICs

Previous studies have shown that SCA sectors overlap in DHFR with residues experimentally determined to be involved in millisecond motions as well as surface residues that allosterically regulate the active site.^25,47,54,55^ These sites include the residues in the Met20 loop. However, there has been little investigation into the mechanisms underlying the allosteric communication in sectors. One possibility is that the motions of positions in the same IC or sector are coupled. To determine how well SCA can predict correlated motions within a protein, we ran a 13 nanosecond (ns) MD simulation on wild-type *E. coli* DHFR (starting structure PDB:3QL3). After extracting the pairwise dynamic cross correlations and constructing a matrix, we were able to determine whether dynamic motions within each IC significantly varied from the rest of the protein **(Figure 2A and 2B)**. Because the ICs are relatively small (8 residues in the case of IC3) and often contain sequential amino acids which can skew correlated motions, the pairwise correlations for residues within 2 positions of each other in the sequence were not considered (i.e. the correlation coefficient between residues 50 and 52 in IC3 was not included in the analysis).

**Figure 2.**
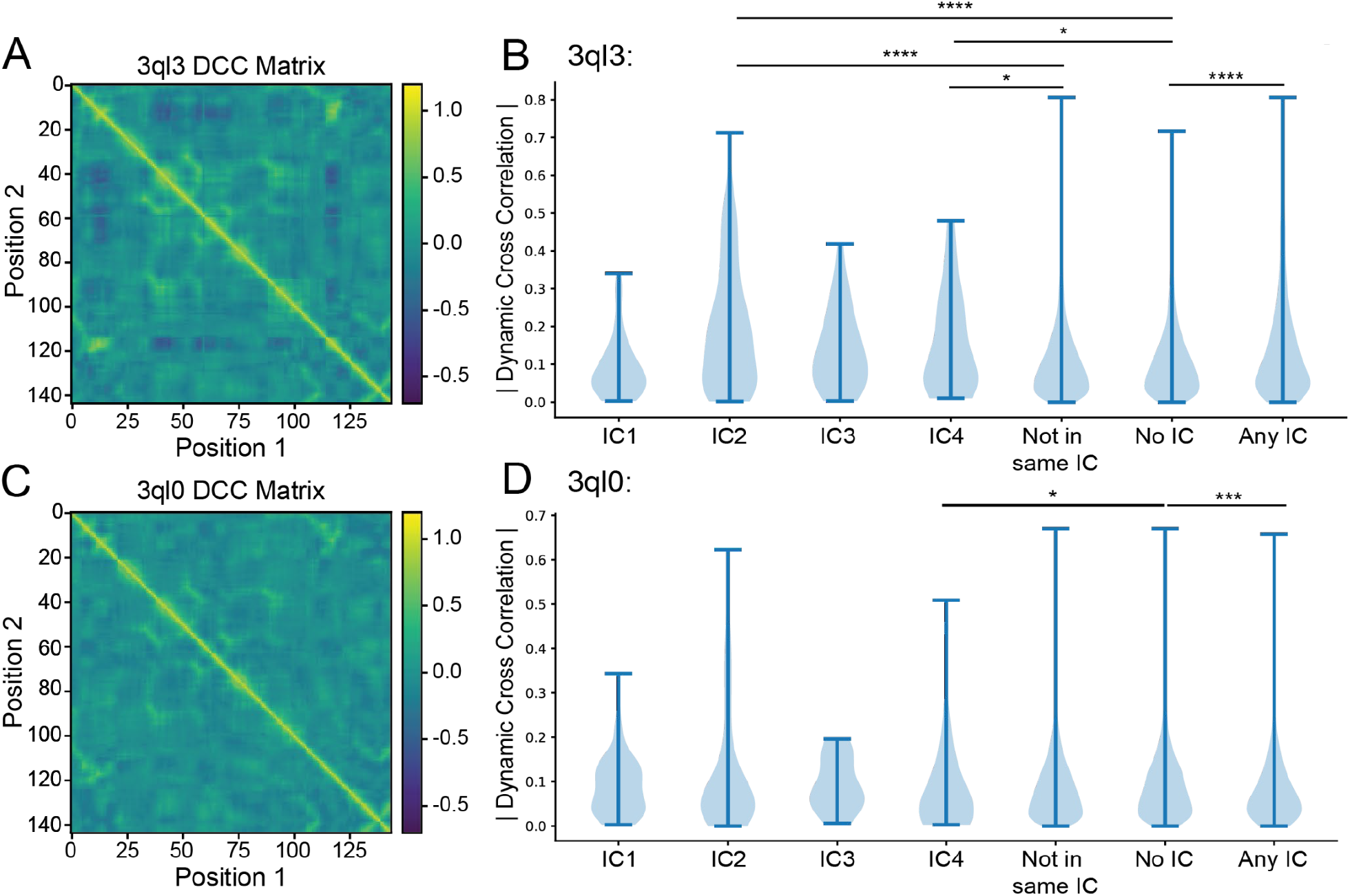
Dynamic cross correlation matrix for wild-type **(A)** and mutant **(C)** *E. coli* DHFR (PDB: 3QL3 and 3QL0, respectively) in the E:NADP+:FOL state and the respective absolute value breakdown of their pairwise motions by independent component **(B and D)**. Categories “Not in Same IC,” “No IC,” and “Any IC” correspond to all pairs of amino acids which lie outside of a given independent component, pairs of amino acids where neither amino acid has an independent component assignment, and all pairs where each amino acid is in assigned to an independent component, respectively. Significance was determined using a two-sided Mann-Whitney U test and labeled following the convention *p≤0.05, **p≤0.01, ***p≤0.001, ****p≤0.0001.

Using a Mann-Whitney U test, it was determined that the correlated motions distributions for IC2 and IC4 were significantly different from those which were measured for residue pairs partially or fully outside of a given IC (“not in same IC”) (p=2.24 * 10^−10^ and p=0.0402, respectively) as well as for those where both residues did not belong to an IC (“no IC”) (p=4.57 *10^−11^ and p=0.0264, respectively) **(Figure 2B)**. IC3, likely due to its small size, did not show significance when compared with the “No IC” assignment nor with “Not in same IC” (p=0.084 and p=0.118, respectively) **(Table S2)**. However, it is apparent from **Figure 2B** that the mean correlated motion of IC3 is higher than that of the “No IC” as well as “Not in same IC” categories. IC1 showed no significant increase in correlated motions. This is in line with the 2015 findings by Teşileanu et al. that the top eigenmode, the mode which represents the most variance, of the SCA matrix may be independent of evolutionary correlations between positions and instead may capture general patterns of conservation.^29^

Interestingly, the distribution of pairwise correlations for all residues assigned to an IC (“Any IC”) were significantly different when compared with “No IC” (p=0.0101). Furthermore, the top pairwise correlations, Gly15-Trp47 and Gly15-Gly121, while captured by SCA, were not captured within any of the ICs and rather occur across multiple ICs. Gly15 belongs to IC2 and Trp47 to IC4 while Gly121 belongs to IC3. This point illustrates the benefit of grouping ICs into sectors especially when considering protein dynamics. Finally, as is apparent by the range of the “No IC” category in **Figure 2B**, SCA ICs do not capture all highly correlated motions. Regardless of mis-grouping top contributors and failing to completely capture all coupled motions, SCA enriched for a disproportionate amount of correlated motions and did so while capturing multiple residues which have been experimentally determined to be catalytically and dynamically important.

### Mutated DHFR Shows Dampened Motions Within ICs

In 2011, the Wright group published a double mutant *E. coli* DHFR, N23PP/S148A, which destabilized the occluded form of the Met20 loop, disrupted millisecond-timescale motions within the active site, and severely reduced catalytic activity.^47^ The authors also created single mutant proteins S148A and N23PP. The S148A mutant trapped the Met20 loop in the closed position but retained millisecond motions in active site residues while the N23PP mutant abrogated active site residue motions. Ser148, located C-terminally from the GH-loop, forms a hydrogen bond with Asn23 which lies directly within the Met20 loop. Disruption of this bond can directly account for the abrogation of Met20 loop conformational changes observed in the generated mutant. However, the mechanism by which the N23PP/S148A double mutant abrogates other active site residue dynamics remains unaccounted for.

To investigate whether disrupting allosteric communication in the ICs of *ec*DHFR can explain the unaccounted for dynamic change, we ran a 13 ns MD simulation on the double mutant. Consistent with the Wright group’s findings, our MD simulation displayed dramatically decreased correlated dynamics relative to wild type. Visual comparison of the dynamic cross correlation matrices in **Figure 2A and 2C** showcases the decreased correlations. After breaking down the dynamic cross correlation matrix by SCA assignment and excluding amino acid pairs within 2 positions of one another, as was done for the wild type, we were able to inspect the role of evolutionarily conserved networks in the abrogated dynamics. Notably, when using a two-sided Mann-Whitney U test, only IC4 retained a significant difference from the “No IC” (p=0.0216). Average values for IC4 and “No IC” are 0.0887 and 0.0943, respectively. Additionally, a one-way Mann-Whitney U test confirmed the “No IC” distribution is greater than IC4 (p=0.0108). Similarly, when examining the difference between “No IC” and “Any IC” (two-sided p=2.70e-04) a one-way Mann-Whitney U revealed that the “No IC” distribution is larger (p=1.35e-04) with the “Any IC” average at 0.0936 **(Table S2)**. Both of these situations are the opposite of what was observed for the wild-type protein.

To further probe the differences between wild-type and mutant dynamics, we performed between groups statistical analysis. Most prominent was the shift for IC2 (p=6.92*10^−8^) and again IC4 (p=3.93*10^−4^) while IC1 and IC3 distributions showed no significant change **(Table S3)**. Again, these findings mirror the initial description of mutant dynamics by the Wright group: dynamics of amino acids around the active site loops are most highly perturbed. The shift for “No IC” and “Any IC” were also both highly significant (p=2.45*10^-12^ and p=6.52*10^−23^, respectively) **(Table S3)**. Overall, these trends indicate that dynamics in IC associated positions, especially IC2 and IC4, were selectively decreased in the N23PP/S148A mutant.

Admittedly, the extent to which our nanosecond timescale MD simulation was able to recapitulate the full scale of dynamic changes in the N23PP/S148A double mutant, which were originally evaluated on the millisecond timescale by Carr-Purcell-Meiboom-Gill (CPMG)–based R2 relaxation dispersion experiments, may be limited.^47^ To probe how well our simulation was able to recapitulate the CPMG results of Michaelis model complex movements, we examined the behavior of three amino acids: Gly121, Gly57, and Ser77. At various points throughout the catalytic cycle, these residues displayed millisecond timescale movements in active site loop conformations, substrate/product binding, and cofactor binding, respectively.^54^ As expected for the Michaelis model complex, we observed the most dramatic changes in active site loop conformation movements (Gly121) while the substrate/product (Gly57) and cofactor binding (Ser77) changes were not as pronounced **(Figure 3)**. Taken together, these results suggest that dampening of an evolutionarily conserved allosteric network within *ec*DHFR, as identified by SCA, serves as the mechanistic link by which the N23PP/ S148A double mutation produces a dramatic knockdown of active site dynamics in *ec*DHFR.

**Figure 3.**
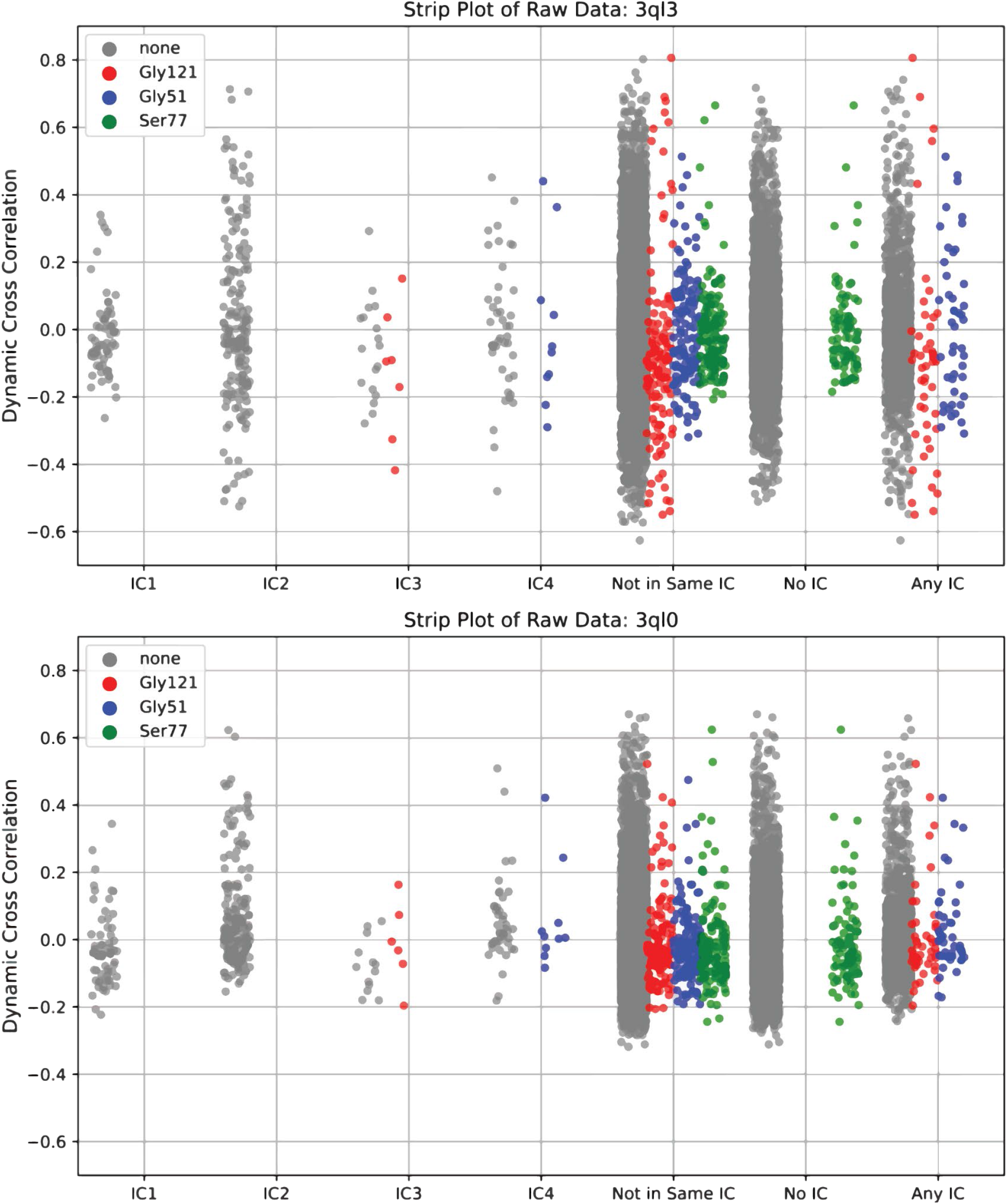
Raw pairwise dynamic cross correlation data for 3QL3 (top) and 3QL0 (bottom). Pairwise correlations involving the residues Gly121, Gly51, and Ser77, respectively labeled red, blue and green, were highlighted for their known millisecond conformational exchange as determined by CPMG and their respective involvement in active site loop conformation, substrate/product binding, and cofactor binding.

### Human DHFR Sector Shows Two Compensatory Mutations

Recent evidence from a group led by Doeke Hekstra advanced the link between Met20 loop dynamics and total *ec*DHFR dynamics. Specifically, the group identified a global hinge motion that is monotonically linked to Met20 loop backbone dihedrals. Interestingly, the authors note that their observed hinge motion is more pronounced in *h*DHFR despite the fact that *h*DHFR bears the N23PP mutation.^61^ In line with the Hekstra group’s findings, our own examination of the N23PP/S148A mutant revealed decreased hinge motions relative to wild-type **(Figure S3A and S3B)**. We have shown that the N23PP/S148A mutant selectively knocks down dynamic communication in an evolutionarily conserved network, and the Hekstra group has established a link between Met20 backbone dihedral dynamics and global hinge motions. How then has human DHFR evolved to retain its dynamic modes while harboring the N23PP mutation?

To probe this question, we first mapped DHFR’s independent components onto *h*DHFR (PDB: 4M6K) **(Table S4)**. Because we previously determined that the top eigenmode (IC1) was not dynamically relevant in *ec*DHFR, we decided to focus our analysis on ICs 2-4, hereafter referred to as a sector. After using PyMOL to map the *E. coli* and human sectors to their respective proteins and create a structure-based alignment **(Figure S4)**, we were able to examine changes to the conserved allosteric network. Two regions in the *h*DHFR sector do not structurally align well with the *E. coli* sector: *h*Pro23/*h*Trp24 and *h*Pro61/*h*Arg65 **(Figure S4B)**. The misalignment at *h*Pro23 and *h*Trp24 relative to *ec*DHFR is unsurprising as these are the two amino acids immediately preceding the Pro-Pro motif from which the N23PP *E. coli* mutation was derived. At the second misaligned site, there is an insertion between the positions *h*Pro61 and *h*Arg65, while the corresponding positions in the *E. coli* sector (*e*Gly51 and *e*Arg52) are contiguous. Interestingly, the *h*61-PEKN-65 sequence has been previously identified to rescue N23PP *E. coli* mutant catalysis when inserted in place of *e*Gly51.^23^ While this catalytic recovery indicates the evolutionary benefit of the *h*61-PEKN-65 in the presence of a Met20 loop that contains a double proline, we are unaware of any direct analysis on dynamics in a G51PEKN N23PP double mutant *ec*DHFR.

After exploring the structural mis-alignments in the *E. coli* and human sectors, we looked to potentially disruptive point mutations in the sector. Using the sector alignment, we determined four places in *h*DHFR that were different: *h*Gly20, *h*Trp113, *h*Phe134, and *h*Ser144. The corresponding positions are *e*Asn18, *e*Met92, *e*Tyr100 and *e*Gly121, respectively. We parsed our original MSA of 4422 DHFR sequences to examine the co-occurrence of these point mutations with the Pro-Pro motif in the Met20 loop. In our MSA, 36 sequences possessed the Met20 Pro-Pro motif. Interestingly, we found that the two instances in which the *h*Gly20 position was a non-glycine residue were the same two instances in which the position corresponding to *h*Pro61 was non-proline **(Table S5)**.

Moreover, glycine occurred in the *h*Gly20 position without the presence of the Met20 Pro-Pro motif in 1858 sequences, and proline occurred in the *h*Pro61 position without the Met20 Pro-Pro motif in 2207 sequences. This indicates the *h*Gly20 and *h*Pro61 mutations likely do not significantly disrupt the function of DHFR which lacks the Met20 Pro-Pro motif. These results along with the previous creation of a catalytically active N23PP/G51PEKN *ec*DHFR mutant strongly indicate that *h*Gly20 and *h*Pro61 mutations are important for maintaining DHFR function in the presence of the Met20 Pro-Pro motif and suggest an evolutionary pathway to sequences with a Met20 Pro-Pro motif.

Finally, in order to contextualize these results in our dynamic analysis, we closely examined the wild-type *ec*DHFR, N23PP/S148A *ec*DHFR, and *h*DHFR structures along with electron densities (PDBs 3QL3, 3QL0, and 4M6K, respectively). In doing so, we noted a dramatic shift in the sidechain rotamer distribution of *e*Ser49 between wild-type and mutant *ec*DHFR. The *e*Ser49 sidechain dynamically communicates between the backbone carbonyl of *e*Asn18 and a water molecule in the wild-type structure (occupancy of 0.5 for each rotamer)^47,62,63^ while the mutant structure shows strong density for a single rotamer of *e*Ser49 that only forms a hydrogen bond with a water **(Figure 4A and 4B)**. This communication is reinforced by the fact that the *e*Asn18 sidechain shows hydrogen bonding with the alpha-helix to which *e*Ser49 belongs. In order to further probe the role of *e*Ser49, we examined the *e*Ser49 sidechain rotamer and Asn18 psi dihedral distributions throughout the stable portion of our wild-type and mutant MD simulations. We found that the *e*Ser49 sidechain rotamer remains static throughout the mutant simulation and shows a distribution of multiple rotamers for the wild type. Additionally, the Asn18 psi dihedral shows biasing towards -130° in the mutant, compared to a more negative dihedral for the wild type, which corresponds to the backbone carbonyl facing farther away from the *e*Ser49 sidechain **(Figure 4)**. As controlled protonation of N5 in Folate is essential to the mechanism of catalysis, perturbations to the *e*Ser49 sidechain rotamer distribution could partially explain the loss of catalytic activity in the N23PP mutant.^56^ However, more experimentation is needed to determine the role of the coordinated water. In the human variant, the structure shows that the *h*Gly20, corresponding to *e*Asn18, backbone carbonyl and *h*Ser59 sidechain, corresponding to *e*Ser49, are not in contact **(Figure 4C)**. Furthermore, *h*Gly20 possesses no sidechain by which to hydrogen bond with the alpha-helix containing *h*Ser59. Based on our analysis and the findings of the Hekstra group, we propose a model whereby the backbone dihedral angles of the Met20 loop in *ec*DHFR allosterically communicate across the active site cleft via *e*Asn18 to effect hinge motion. To escape the deleterious effect that the Met20 loop Pro-Pro motif would have on the Met20 loop dihedrals and consequently overall hinge motion, the human sector has evolved a Glycine at the *e*Asn18 position. In depth experimentation would be needed to confirm this model.

**Figure 4.**
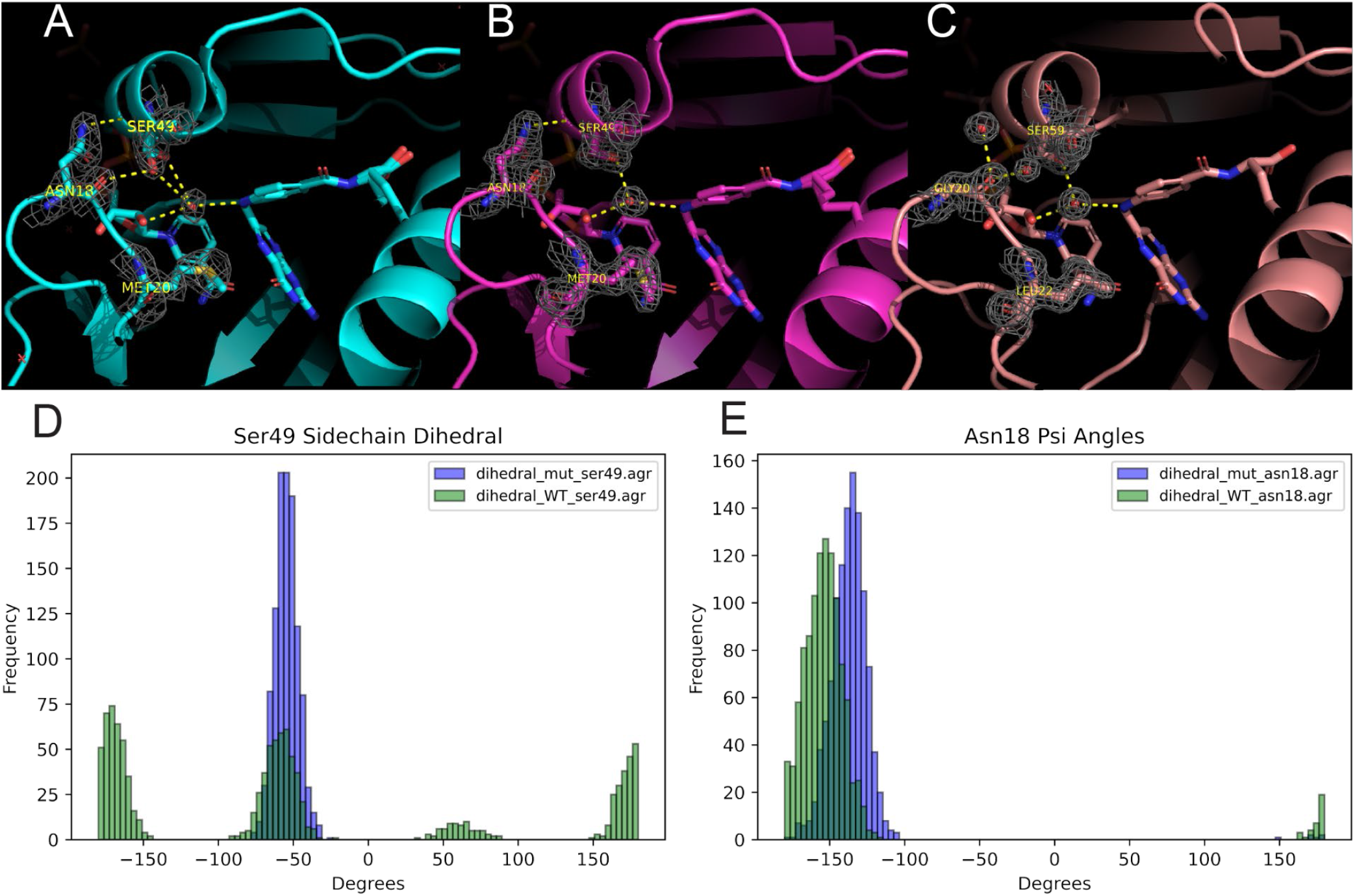
PDB structures for wild-type *ec*DHFR **(A)**, N23PP/S148A mutant *ec*DHFR **(B)**, and *h*DHFR **(C)**, with electron density (3QL3, 3QL0, and 4M6K, respectively). Dihedral distribution for the Ser49 sidechain **(D)** and Asn18 Psi angle **(E)** throughout wild-type (green) and mutant (purple) MD simulations assessed for frames 1500-2600.

## Conclusion

This paper is the first example to demonstrate the ability of SCA to identify networks of coevolving residues in an enzyme which exhibit an increase in correlated motions relative to the remainder of the protein. We demonstrated that increased correlation in dynamics are apparent when viewing these networks from both the individual IC level as well as grouping ICs into a sector. Furthermore, we were able to show that the dynamic communication of this network was selectively decreased in a previously designed mutant *ec*DHFR dynamic knockout. Specifically, the relative change of “Any IC,” IC2, and IC4 with respect to “No IC” shows a clear selective decrease in the dynamics of SCA-identified residues over non-IC residues. Through the lens of SCA, we also gained insight into how conserved networks within DHFR have coevolved to accommodate an otherwise dynamically deleterious mutation present in human DHFR. We believe that our model of allosteric communication from the Met20 loop across *e*Asn18 to *e*Ser49 offers an explanation and starting point for future experimentation as to how human DHFR has at least partially decoupled Met20 backbone dihedrals from global hinge motion. More broadly, our findings implicate protein dynamics as a driving force for evolution. That is, our work has shown that electronic perturbations to networks of amino acids which exhibit correlated dynamics are more likely to be evolutionarily selected against. This concept will undoubtedly prove useful in the design and understanding of enzymes.

## Acknowledgements

We would like to thank the National Institutes of Health for financial support for this work, specifically CMFA, TLK, JMM, ASO, and ASW acknowledge R35GM146987 and CIC acknowledges 5T32GM008320-34 for support. The content is solely the responsibility of the authors and does not necessarily represent the official views of the National Institutes of Health. We would also like to thank the Vanderbilt Center for Structural Biology, Dr. Jarrod Smith, and the Vanderbilt ACCRE core for computational resources.

## Notes

### Competing Interest Statement

The authors have declared no competing interest.

## References

(1) Kamerlin, S. C. L.; Warshel, A. At the Dawn of the 21st Century: Is Dynamics the Missing Link for Understanding Enzyme Catalysis? Proteins 2010, 78 (6), 1339–1375.

(2) Kohen, A. Role of Dynamics in Enzyme Catalysis: Substantial versus Semantic Controversies. Acc. Chem. Res. 2015, 48 (2), 466–473.

(3) Jindal, G.; Warshel, A. Misunderstanding the Preorganization Concept Can Lead to Confusions about the Origin of Enzyme Catalysis. Proteins 2017, 85 (12), 2157–2161.

(4) Schwartz, S. D. Protein Dynamics and Enzymatic Catalysis. J. Phys. Chem. B 2023, 127 (12), 2649–2660.

(5) Hammes-Schiffer, S.; Benkovic, S. J. Relating Protein Motion to Catalysis. Annu. Rev. Biochem. 2006, 75, 519–541.

(6) Nam, K.; Wolf-Watz, M. Protein Dynamics: The Future Is Bright and Complicated! Struct Dyn 2023, 10 (1), 014301.

(7) Abali, E. E.; Skacel, N. E.; Celikkaya, H.; Hsieh, Y.-C. Regulation of Human Dihydrofolate Reductase Activity and Expression. Vitam. Horm. 2008, 79, 267–292.

(8) Fierke, C. A.; Johnson, K. A.; Benkovic, S. J. Construction and Evaluation of the Kinetic Scheme Associated with Dihydrofolate Reductase from Escherichia Coli. Biochemistry 1987, 26 (13), 4085–4092.

(9) Li, J.; Lin, J.; Kohen, A.; Singh, P.; Francis, K.; Cheatum, C. M. Evolution of Optimized Hydride Transfer Reaction and Overall Enzyme Turnover in Human Dihydrofolate Reductase. Biochemistry 2021, 60 (50), 3822–3828.

(10) Schnell, J. R.; Dyson, H. J.; Wright, P. E. Structure, Dynamics, and Catalytic Function of Dihydrofolate Reductase. Annu. Rev. Biophys. Biomol. Struct. 2004, 33, 119–140.

(11) Blount, B. C.; Mack, M. M.; Wehr, C. M.; MacGregor, J. T.; Hiatt, R. A.; Wang, G.; Wickramasinghe, S. N.; Everson, R. B.; Ames, B. N. Folate Deficiency Causes Uracil Misincorporation into Human DNA and Chromosome Breakage: Implications for Cancer and Neuronal Damage. Proc. Natl. Acad. Sci. U. S. A. 1997, 94 (7), 3290–3295.

(12) Huennekens, F. M. Folic Acid Coenzymes in the Biosynthesis of Purines and Pyrimidines. In Vitamins & Hormones; Harris, R. S., Wool, I. G., Loraine, J. A., Thimann, K. V., Eds.; Academic Press, 1969; Vol. 26, pp 375–394.

(13) Galassi, R.; Oumarou, C. S.; Burini, A.; Dolmella, A.; Micozzi, D.; Vincenzetti, S.; Pucciarelli, S. A Study on the Inhibition of Dihydrofolate Reductase (DHFR) from Escherichia Coli by Gold(i) Phosphane Compounds. X-Ray Crystal Structures of (4,5-Dichloro-1H-Imidazolate-1-Yl)-Triphenylphosphane-Gold(i) and (4,5-Dicyano-1H-Imidazolate-1-Yl)-Triphenylphosphane-Gold(i). Dalton Trans. 2015, 44 (7), 3043–3056.

(14) Schweitzer, B. I.; Dicker, A. P.; Bertino, J. R. Dihydrofolate Reductase as a Therapeutic Target. FASEB J. 1990, 4 (8), 2441–2452.

(15) Sharma, M.; Chauhan, P. M. S. Dihydrofolate Reductase as a Therapeutic Target for Infectious Diseases: Opportunities and Challenges. Future Med. Chem. 2012, 4 (10), 1335–1365.

(16) Raimondi, M. V.; Randazzo, O.; La Franca, M.; Barone, G.; Vignoni, E.; Rossi, D.; Collina, S. DHFR Inhibitors: Reading the Past for Discovering Novel Anticancer Agents. Molecules 2019, 24 (6). 10.3390/molecules24061140.

(17) Srinivasan, B.; Tonddast-Navaei, S.; Roy, A.; Zhou, H.; Skolnick, J. Chemical Space of Escherichia Coli Dihydrofolate Reductase Inhibitors: New Approaches for Discovering Novel Drugs for Old Bugs. Med. Res. Rev. 2019, 39 (2), 684–705.

(18) Bhabha, G.; Ekiert, D. C.; Jennewein, M.; Zmasek, C. M.; Tuttle, L. M.; Kroon, G.; Dyson, H. J.; Godzik, A.; Wilson, I. A.; Wright, P. E. Divergent Evolution of Protein Conformational Dynamics in Dihydrofolate Reductase. Nat. Struct. Mol. Biol. 2013, 20 (11), 1243–1249.

(19) Tuttle, L. M.; Dyson, H. J.; Wright, P. E. Side Chain Conformational Averaging in Human Dihydrofolate Reductase. Biochemistry 2014, 53 (7), 1134–1145.

(20) Wallace, L. A.; Robert Matthews, C. Highly Divergent Dihydrofolate Reductases Conserve Complex Folding Mechanisms. J. Mol. Biol. 2002, 315 (2), 193–211.

(21) Reddish, M. J.; Vaughn, M. B.; Fu, R.; Dyer, R. B. Ligand-Dependent Conformational Dynamics of Dihydrofolate Reductase. Biochemistry 2016, 55 (10), 1485–1493.

(22) Goldstein, M.; Goodey, N. M. Distal Regions Regulate Dihydrofolate Reductase-Ligand Interactions. Methods Mol. Biol. 2021, 2253, 185–219.

(23) Liu, C. T.; Hanoian, P.; French, J. B.; Pringle, T. H.; Hammes-Schiffer, S.; Benkovic, S. J. Functional Significance of Evolving Protein Sequence in Dihydrofolate Reductase from Bacteria to Humans. Proc. Natl. Acad. Sci. U. S. A. 2013, 110 (25), 10159–10164.

(24) Miller, G. P.; Benkovic, S. J. Stretching Exercises--Flexibility in Dihydrofolate Reductase Catalysis. Chem. Biol. 1998, 5 (5), R105–R113.

(25) Reynolds, K. A.; McLaughlin, R. N.; Ranganathan, R. Hot Spots for Allosteric Regulation on Protein Surfaces. Cell 2011, 147 (7), 1564–1575.

(26) Dokholyan, N. V. Controlling Allosteric Networks in Proteins. Chem. Rev. 2016, 116 (11), 6463–6487.

(27) Perica, T.; Marsh, J. A.; Sousa, F. L.; Natan, E.; Colwell, L. J.; Ahnert, S. E.; Teichmann, S. The Emergence of Protein Complexes: Quaternary Structure, Dynamics and Allostery. Colworth Medal Lecture. Biochem. Soc. Trans. 2012, 40 (3), 475–491.

(28) Halabi, N.; Rivoire, O.; Leibler, S.; Ranganathan, R. Protein Sectors: Evolutionary Units of Three-Dimensional Structure. Cell 2009, 138 (4), 774–786.

(29) Teşileanu, T.; Colwell, L. J.; Leibler, S. Protein Sectors: Statistical Coupling Analysis versus Conservation. PLoS Comput. Biol. 2015, 11 (2), e1004091.

(30) Rajasekaran, N.; Naganathan, A. N. A Self-Consistent Structural Perturbation Approach for Determining the Magnitude and Extent of Allosteric Coupling in Proteins. Biochem. J 2017, 474 (14), 2379–2388.

(31) Lakhani, B.; Thayer, K. M.; Hingorani, M. M.; Beveridge, D. L. Evolutionary Covariance Combined with Molecular Dynamics Predicts a Framework for Allostery in the MutS DNA Mismatch Repair Protein. J. Phys. Chem. B 2017, 121 (9), 2049–2061.

(32) Lakhani, B.; Thayer, K. M.; Black, E.; Beveridge, D. L. Spectral Analysis of Molecular Dynamics Simulations on PDZ: MD Sectors. J. Biomol. Struct. Dyn. 2020, 38 (3), 781–790.

(33) Lee, J.; Natarajan, M.; Nashine, V. C.; Socolich, M.; Vo, T.; Russ, W. P.; Benkovic, S. J.; Ranganathan, R. Surface Sites for Engineering Allosteric Control in Proteins. Science 2008, 322 (5900), 438–442.

(34) McCormick, J. W.; Pincus, D.; Resnekov, O.; Reynolds, K. A. Strategies for Engineering and Rewiring Kinase Regulation. Trends Biochem. Sci. 2020, 45 (3), 259–271.

(35) Pincus, D.; Pandey, J. P.; Feder, Z. A.; Creixell, P.; Resnekov, O.; Reynolds, K. A. Engineering Allosteric Regulation in Protein Kinases. Sci. Signal. 2018, 11 (555). 10.1126/scisignal.aar3250.

(36) Rosensweig, C.; Reynolds, K. A.; Gao, P.; Laothamatas, I.; Shan, Y.; Ranganathan, R.; Takahashi, J. S.; Green, C. B. An Evolutionary Hotspot Defines Functional Differences between CRYPTOCHROMES. Nat. Commun. 2018, 9 (1), 1138.

(37) Rivoire, O.; Reynolds, K. A.; Ranganathan, R. Evolution-Based Functional Decomposition of Proteins. PLoS Comput. Biol. 2016, 12 (6), e1004817.

(38) Narayanan, C.; Gagné, D.; Reynolds, K. A.; Doucet, N. Conserved Amino Acid Networks Modulate Discrete Functional Properties in an Enzyme Superfamily. Sci. Rep. 2017, 7 (1), 3207.

(39) Salinas, V. H.; Ranganathan, R. Coevolution-Based Inference of Amino Acid Interactions Underlying Protein Function. Elife 2018, 7. 10.7554/eLife.34300.

(40) del Sol, A.; Tsai, C.-J.; Ma, B.; Nussinov, R. The Origin of Allosteric Functional Modulation: Multiple Pre-Existing Pathways. Structure 2009, 17 (8), 1042–1050.

(41) Ribeiro, A. A. S. T.; Ortiz, V. A Chemical Perspective on Allostery. Chem. Rev. 2016, 116 (11), 6488–6502.

(42) Wu, N.; Barahona, M.; Yaliraki, S. N. Allosteric Communication and Signal Transduction in Proteins. Curr. Opin. Struct. Biol. 2024, 84, 102737.

(43) Singh, S.; Mandlik, V.; Shinde, S. Molecular Dynamics Simulations and Statistical Coupling Analysis of GPI12 in L. Major: Functional Co-Evolution and Conservedness Reveals Potential Drug-Target Sites. Mol. Biosyst. 2015, 11 (3), 958–968.

(44) Case, D. A.; Metin Aktulga, H.; Belfon, K.; Ben-Shalom, I.; Berryman, J. T.; Brozell, S. R.; Cerutti, D. S.; Cheatham, T. E., III; Andrés Cisneros, G.; Cruzeiro, V. W. D.; Darden, T. A.; Duke, R. E.; Giambasu, G.; Gilson, M. K.; Gohlke, H.; Goetz, A. W.; Harris, R.; Izadi, S.; Izmailov, S. A.; Kasavajhala, K.; Kaymak, M. C.; King, E.; Kovalenko, A.; Kurtzman, T.; Lee, T.; LeGrand, S.; Li, P.; Lin, C.; Liu, J.; Luchko, T.; Luo, R.; Machado, M.; Man, V.; Manathunga, M.; Merz, K. M.; Miao, Y.; Mikhailovskii, O.; Monard, G.; Nguyen, H.; O’Hearn, K. A.; Onufriev, A.; Pan, F.; Pantano, S.; Qi, R.; Rahnamoun, A.; Roe, D. R.; Roitberg, A.; Sagui, C.; Schott-Verdugo, S.; Shajan, A.; Shen, J.; Simmerling, C. L.; Skrynnikov, N. R.; Smith, J.; Swails, J.; Walker, R. C.; Wang, J.; Wang, J.; Wei, H.; Wolf, R. M.; Wu, X.; Xiong, Y.; Xue, Y.; York, D. M.; Zhao, S.; Kollman, P. A. Amber 2022; University of California, San Francisco, 2022.

(45) Case, D. A.; Aktulga, H. M.; Belfon, K.; Cerutti, D. S.; Cisneros, G. A.; Cruzeiro, V. W. D.; Forouzesh, N.; Giese, T. J.; Götz, A. W.; Gohlke, H.; Izadi, S.; Kasavajhala, K.; Kaymak, M. C.; King, E.; Kurtzman, T.; Lee, T.-S.; Li, P.; Liu, J.; Luchko, T.; Luo, R.; Manathunga, M.; Machado, M. R.; Nguyen, H. M.; O’Hearn, K. A.; Onufriev, A. V.; Pan, F.; Pantano, S.; Qi, R.; Rahnamoun, A.; Risheh, A.; Schott-Verdugo, S.; Shajan, A.; Swails, J.; Wang, J.; Wei, H.; Wu, X.; Wu, Y.; Zhang, S.; Zhao, S.; Zhu, Q.; Cheatham, T. E., 3rd; Roe, D. R.; Roitberg, A.; Simmerling, C.; York, D. M.; Nagan, M. C.; Merz, K. M., Jr. AmberTools. J. Chem. Inf. Model. 2023, 63 (20), 6183–6191.

(46) Berman, H.; Henrick, K.; Nakamura, H. Announcing the Worldwide Protein Data Bank. Nat. Struct. Biol. 2003, 10 (12), 980.

(47) Bhabha, G.; Lee, J.; Ekiert, D. C.; Gam, J.; Wilson, I. A.; Dyson, H. J.; Benkovic, S. J.; Wright, P. E. A Dynamic Knockout Reveals That Conformational Fluctuations Influence the Chemical Step of Enzyme Catalysis. Science 2011, 332 (6026), 234–238.

(48) H++ (web-based computational prediction of protonation states and pK of ionizable groups in macromolecules). http://biophysics.cs.vt.edu/H++ (accessed 2024-04-19).

(49) Tian, C.; Kasavajhala, K.; Belfon, K. A. A.; Raguette, L.; Huang, H.; Migues, A. N.; Bickel, J.; Wang, Y.; Pincay, J.; Wu, Q.; Simmerling, C. ff19SB: Amino-Acid-Specific Protein Backbone Parameters Trained against Quantum Mechanics Energy Surfaces in Solution. J. Chem. Theory Comput. 2020, 16 (1), 528–552.

(50) He, X.; Man, V. H.; Yang, W.; Lee, T.-S.; Wang, J. A Fast and High-Quality Charge Model for the next Generation General AMBER Force Field. J. Chem. Phys. 2020, 153 (11), 114502.

(51) Roe, D. R.; Cheatham, T. E., 3rd. PTRAJ and CPPTRAJ: Software for Processing and Analysis of Molecular Dynamics Trajectory Data. J. Chem. Theory Comput. 2013, 9 (7), 3084–3095.

(52) Website. Schrödinger, L. & DeLano, W., 2020. PyMOL, Available at: http://www.pymol.org/pymol.

(53) Chen, J.; Dima, R. I.; Thirumalai, D. Allosteric Communication in Dihydrofolate Reductase: Signaling Network and Pathways for Closed to Occluded Transition and Back. J. Mol. Biol. 2007, 374 (1), 250–266.

(54) Boehr, D. D.; McElheny, D.; Dyson, H. J.; Wright, P. E. The Dynamic Energy Landscape of Dihydrofolate Reductase Catalysis. Science 2006, 313 (5793), 1638–1642.

(55) McElheny, D.; Schnell, J. R.; Lansing, J. C.; Dyson, H. J.; Wright, P. E. Defining the Role of Active-Site Loop Fluctuations in Dihydrofolate Reductase Catalysis. Proc. Natl. Acad. Sci. U. S. A. 2005, 102 (14), 5032–5037.

(56) Liu, C. T.; Francis, K.; Layfield, J. P.; Huang, X.; Hammes-Schiffer, S.; Kohen, A.; Benkovic, S. J. Escherichia Coli Dihydrofolate Reductase Catalyzed Proton and Hydride Transfers: Temporal Order and the Roles of Asp27 and Tyr100. Proc. Natl. Acad. Sci. U. S. 2014, 111 (51), 18231–18236.

(57) Singh, P.; Sen, A.; Francis, K.; Kohen, A. Extension and Limits of the Network of Coupled Motions Correlated to Hydride Transfer in Dihydrofolate Reductase. J. Am. Chem. Soc. 2014, 136 (6), 2575–2582.

(58) Francis, K.; Kohen, A. Protein Motions and the Activation of the CH Bond Catalyzed by Dihydrofolate Reductase. Curr. Opin. Chem. Biol. 2014, 21, 19–24.

(59) Liu, C. T.; Layfield, J. P.; Stewart, R. J., 3rd; French, J. B.; Hanoian, P.; Asbury, J. B.; Hammes-Schiffer, S.; Benkovic, S. J. Probing the Electrostatics of Active Site Microenvironments along the Catalytic Cycle for Escherichia Coli Dihydrofolate Reductase. J. Am. Chem. Soc. 2014, 136 (29), 10349–10360.

(60) Mhashal, A. R.; Pshetitsky, Y.; Eitan, R.; Cheatum, C. M.; Kohen, A.; Major, D. T. Effect of Asp122 Mutation on the Hydride Transfer in E. Coli DHFR Demonstrates the Goldilocks of Enzyme Flexibility. J. Phys. Chem. B 2018, 122 (33), 8006–8017.

(61) Greisman, J. B.; Dalton, K. M.; Brookner, D. E.; Klureza, M. A.; Sheehan, C. J.; Kim, I.-S.; Henning, R. W.; Russi, S.; Hekstra, D. R. Perturbative Diffraction Methods Resolve a Conformational Switch That Facilitates a Two-Step Enzymatic Mechanism. Proc. Natl. Acad. Sci. U. S. A. 2024, 121 (9), e2313192121.

(62) Sehnal, D.; Bittrich, S.; Deshpande, M.; Svobodová, R.; Berka, K.; Bazgier, V.; Velankar, S.; Burley, S. K.; Koča, J.; Rose, A. S. Mol* Viewer: Modern Web App for 3D Visualization and Analysis of Large Biomolecular Structures. Nucleic Acids Res. 2021, 49 (W1), W431–W437.

(63) Berman, H. M.; Westbrook, J.; Feng, Z.; Gilliland, G.; Bhat, T. N.; Weissig, H.; Shindyalov, I. N.; Bourne, P. E. The Protein Data Bank. Nucleic Acids Res. 2000, 28 (1), 235–242.

