## Supplemental Information for "Statistical Coupling Analysis Predicts Correlated Motions in Dihydrofolate Reductase"

### Supporting Information

**Table S1.** *E. coli* IC residue compositions at cutoff = 0.95.

|  | *ec*DHFR residues |
| --- | --- |
| IC1 | 13,23,25,27,32,39,55,63,71,107,133,153 |
| IC2 | 7,14,15,31,35,42,43,44,46,49,53,54,57,59,94,95,96,100,113,122,126 |
| IC3 | 21,22,24,50,52,64,81,121, |
| IC4 | 5,6,11,18,40,45,47,51,92,111,125, |

**Table S2.** Within variant p-values from Mann-Whitney U tests of dynamic correlations.

| **Comparison Całegory** | **3QL3** | **3QLO** |
| --- | --- | --- |
| IC1 vs No IC | 0.11780 | 0.74632 |
| IC2 vs No IC | 4.57248E-11 | 0.39920 |
| IC3 vs No IC | 0.12111 | 0.59298 |
| IC4 vs No IC | 0.02640 | 0.02156 |
| IC1 vs Not in Same IC | 0.08413 | 0.39950 |
| IC2 vs Not in Same IC | 2.23752E-10 | 0.89636 |
| IC3 vs Not in Same IC | 0.15424 | 0.41011 |
| IC4 vs Not in Same IC | 0.04020 | 0.05310 |
| Any IC vs No IC | 8.03962E-08 | 2.69767E-04 |
| Not in Same IC vs No IC | 0.25586 | 5.37196E-04 |

**Table S3.** Across variant p-values from Mann-Whitney U tests of dynamic correlations.

| **Comparison Category** | **Comparison** |
| --- | --- |
| IC1 (3QL3) vs IC1 (3QL0) | 0.6576 |
| IC2 (3QL3) vs IC2 (3QL0) | 6.92E-08 |
| IC3 (3QL3) vs IC3 (3QL0) | 0.24081 |
| IC4 (3QL3) vs IC4 (3QL0) | 3.93E-04 |
| Any IC (3QL3) vs Any IC (3QL0) | 6.52E-23 |
| No IC (3QL3) vs No IC (3QL0) | 2.45E-12 |
| Not in Same IC (3QL3) vs Not in Same IC (3QL0) | 4.62E-58 |

**Table S4.** Human DHFR IC residue compositions at cutoff = 0.95.

|  | Human DHFR residues |
| --- | --- |
| IC1 | 15,26,28,30,35,47,49,68,76,85,128,156,179 |
| IC2 | 9,16,17,34,38,52,53,54,56,59,66,67,70,72,115,116,117,121,136,145,149 |
| IC3 | 23,24,27,60,65,77,96,144 |
| IC4 | 7,8,13,20,50,55,57,61,113,134,148 |

**TableS5.** Proteins for which the Met20 Pro-Pro motif was present in our original alignment and which did not possess either the Gly20 mutation or the exact 61-PEKN-65 mutation. PFAM protein accession numbers are in the first column with common or scientific names available in the second column.

| **PFAM accession** | **Name** | **Gly20** | **Met20**  **Loop motif** | **P61-N65** |
| --- | --- | --- | --- | --- |
| R7UI73 | *Capitella teleta*  (Polychaete worm) | C | PP | GEEE |
| U6GUT1 | *Eimeria acervulina*  (Coccidian parasite) | N | PP | empty |
| S7NYA8 | brandts bat | G | PP | PKKN |
| G1QES9 | little brown bt | G | PP | PKKN |
| L5KJI5 | black flying fox | G | PP | PKKN |
| S7NHW6 | brandts bat | G | PP | PKKN |
| M1VWK3 | *claviceps purpurea* (ergot) | G | PP | PPSF |
| A0A2Z5U771 | *Rhinolophus gammaherpesvirus 1* (from greater horshoe bat) | G | PP | PTKS |
| A0A3L8S6E8 | *Chloebia gouldiae*  (Gouldian finch) | G | PP | PEKS |
| U3IIA3 | *Anas platyrhynchos platyrhynchos*  (Northern mallard) | G | PP | PEKH |

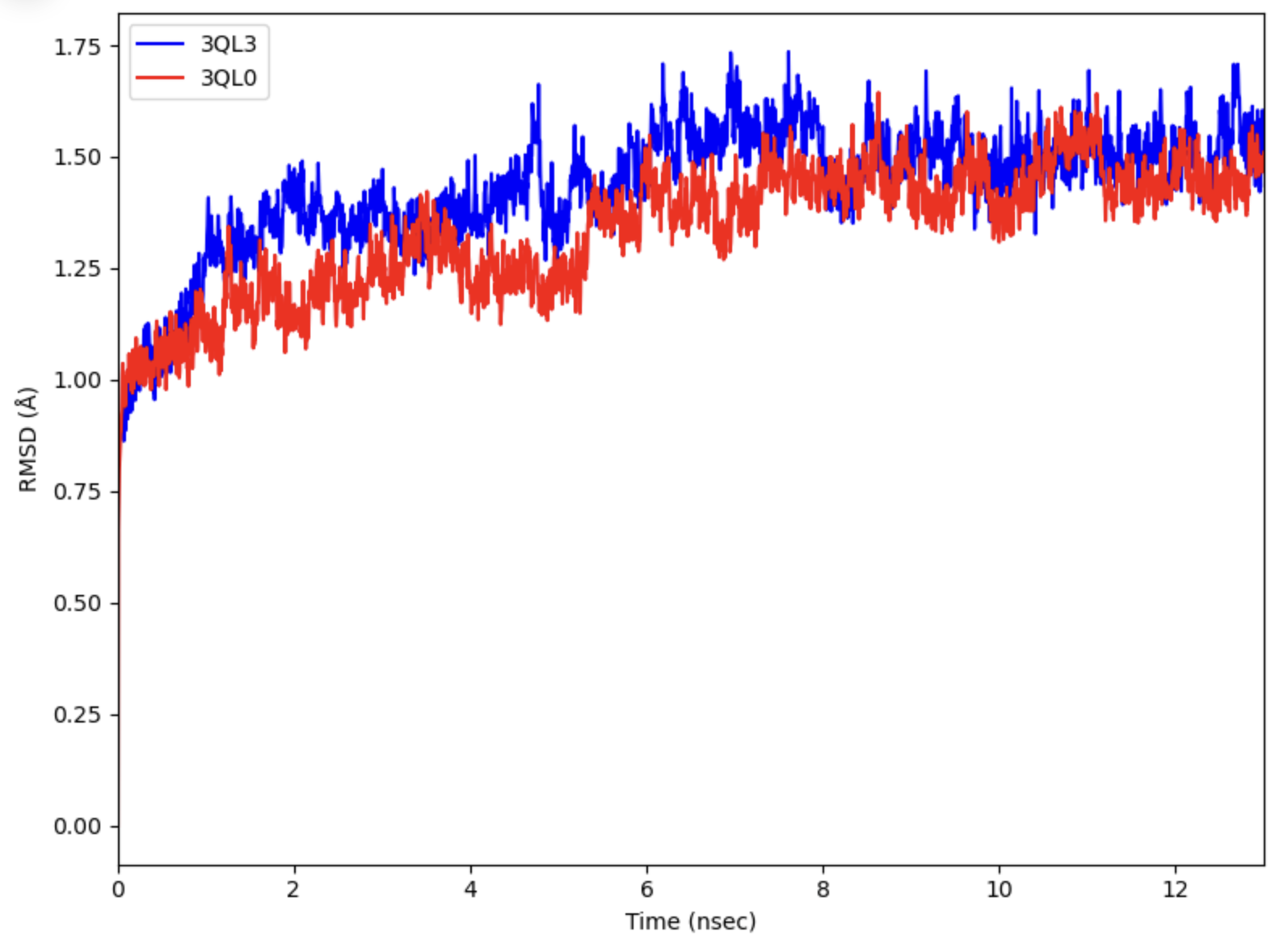

**Figure S1.** The root-mean squared deviation (RMSD) of wild-type (3QL3, blue) and mutant (3QL0, red) DHFR with respect to the initial structure over the duration of 13 ns simulation.

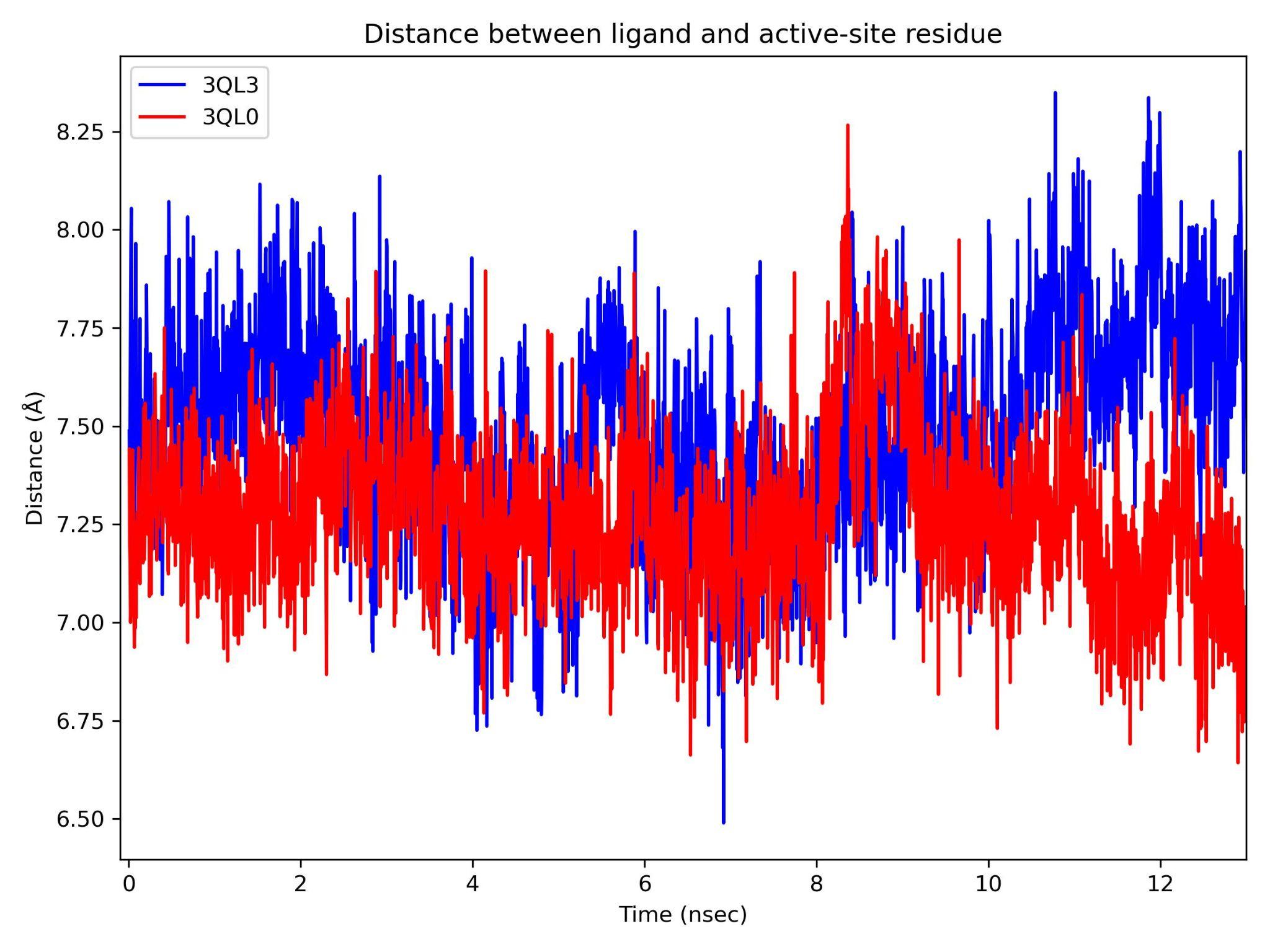

**Figure S2.** The distance between the folate ligand and Asp residue in the active site of wild-type (blue) and mutant (red) DHFR over the course of simulation.

**Simulation stability**.

Figure S1 monitors the RMSD of the proteins with respect to the first frame of the trajectory over the simulation time. Simulations of the wild-type and mutant DHFR are shown to be reasonably stable, with fluctuations gradually increasing then stabilizing at approximately 1.5 Å starting at around 6 ns. Distance between the folate ligand and a residue in the DHFR active site (D27 in 3QL3 and D28 in 3QL0) from center of mass was also calculated to monitor the stability of the ligand-amino acid interaction during the simulation. Figure S2 shows this distance averages at around 7.4 Å and fluctuations are within a range of 2 Å for both the wild-type and mutant DHFR, suggesting relative stability of this interaction throughout the simulation.

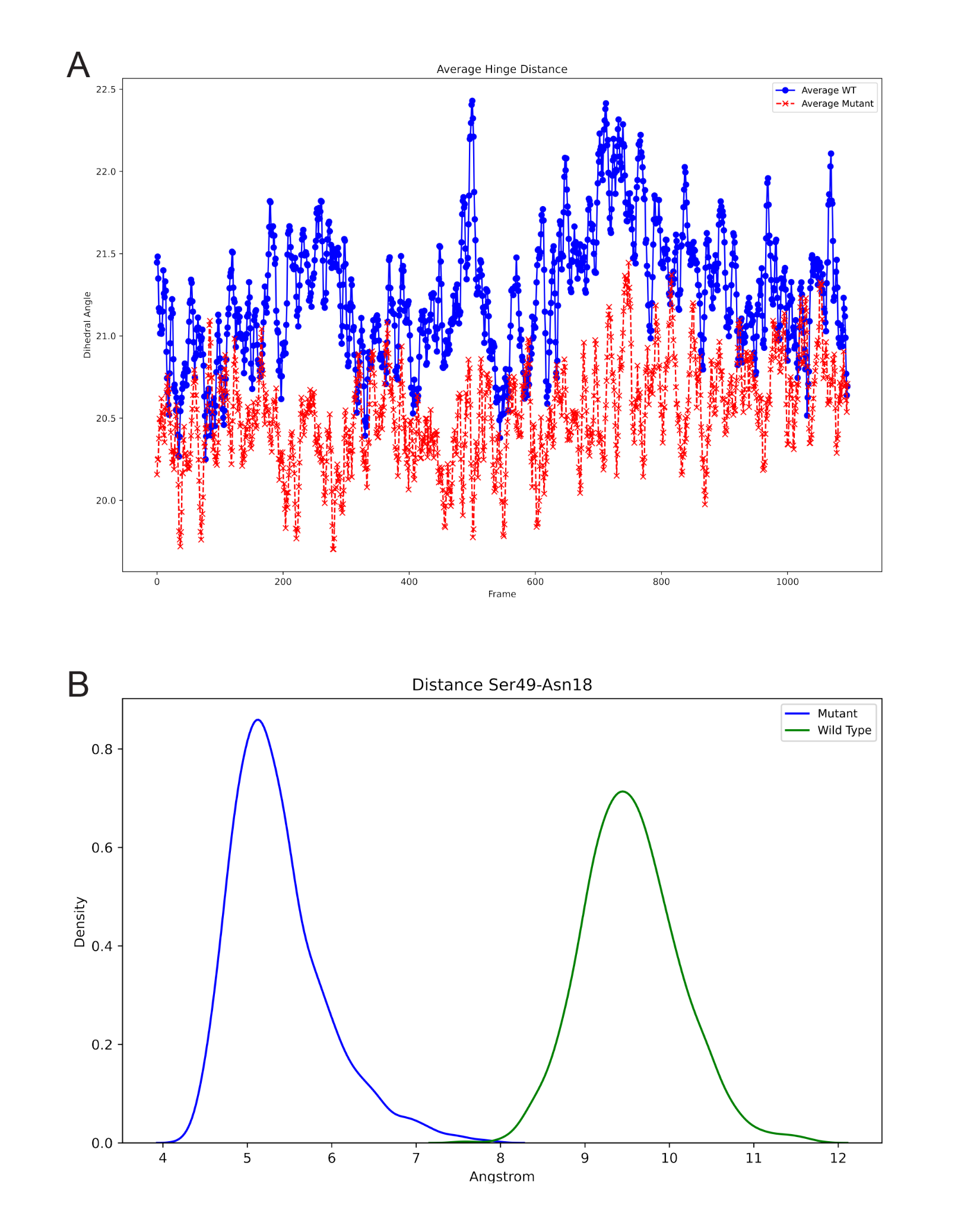

**Figure S3. (A)** shows the average hinge distance for wild-type and mutant DHFR. Wild-type hinge distance (blue) is the average distance between positions 22-53, 23-53 and 24-53 while mutant hinge distance (red) is the average of the distance between positions 22-54, 23-54 and 25-54. **(B)** shows the kernel density estimate of the distance between Ser49 and Asn18 in wild-type (green) and mutant (blue). All measurements were taken at the alpha carbon and measured for simulation frames 1500-2600.

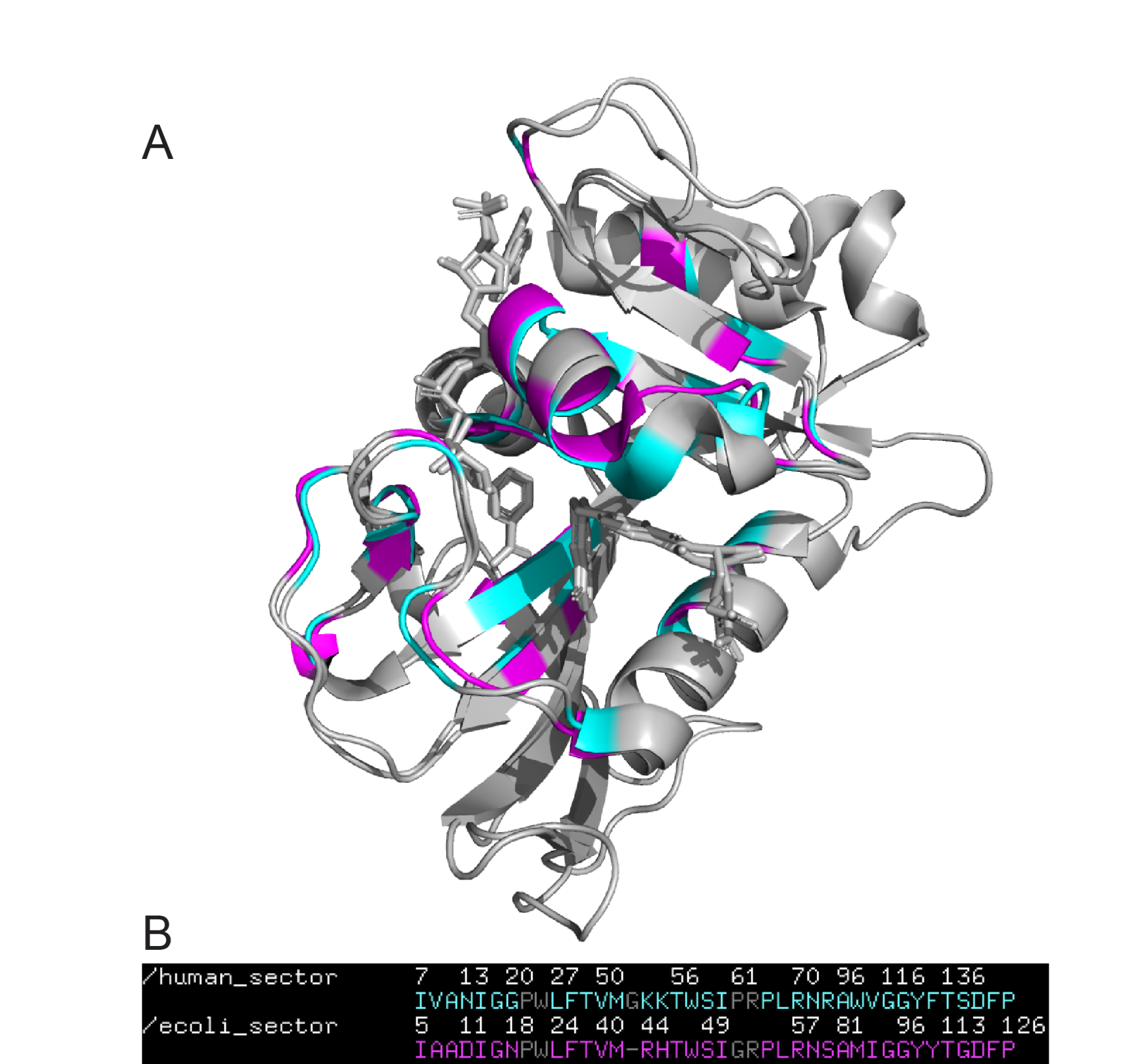

**Figure S4.** Overlaid PDB structures for human (PDB:4M6K) and *E. coli* (PDB:3QL3) DHFR **(A)** along with the structure-based alignment of their sectors (ICs 2-4) **(B)**. The RMSD for the structure-based alignment was 0.853Å after 5 cycles with a total of 42 rejected atoms. Numbering is only accurate for the amino acid over which it appears and is meant as a reference for the others.
